# Dysregulated Protein Translation Control in Hidradenitis Suppurativa: Implication for Lesion-associated Squamous Cell Carcinoma Development

**DOI:** 10.1101/2021.10.14.464300

**Authors:** Lin Jin, Mahendra P. Kashyap, Yunjia Chen, Yuanyuan Guo, Jasim Khan, Jari Q. Chen, Madison B Lee, Zhiping Weng, Allen Oak, Rajesh Sinha, M. Shahid Mukhtar, Jessy S. Deshane, Chander Raman, Craig A. Elmets, Mohammad Athar

**Affiliations:** UAB Research Center of Excellence in Arsenicals, Department of Dermatology, School of Medicine, University of Alabama at Birmingham, AL 35294, USA; UAB Medical Genomics Laboratory, Department of Genetics, University of Alabama at Birmingham, AL 35294, USA; Hoover High School, Hoover, AL, 35244, USA; School of Medicine, University of Alabama at Birmingham, AL 35294, USA; Department of Biology, University of Alabama at Birmingham, Birmingham, AL 35294, USA; Division of Pulmonary, Allergy and Critical Care Medicine, University of Alabama at Birmingham, AL 35294, USA

## Abstract

Hidradenitis Suppurativa (HS) is a proinflammatory painful skin disorder. This chronic disease is often associated with aggressive squamous cell carcinoma (SCC). The molecular pathogenesis of this disease progression remains undefined. The translation initiation factor eIF4E/4G/4A1 complex is overexpressed in a variety of human malignancies. In this study, we found that the expression of eIF4E/4G/4A1 as well as phosphorylated eIF4E were upregulated in HS skin. In the global transcription profiles derived from two public database, we were able to enrich 734 eIF4F-related genes. GSEA pathway enrichment analysis further demonstrated that RAS/MEK/ERK oncogene signaling pathway associated with inflammation signaling were significantly activated in HS lesion. The increase expression of eIF4 protein components was associated with enhanced eIF4E translation targets Cyclin D1 and c-Myc. Confocal fluorescence microscopy analysis further revealed that Cyclin D1 and c-Myc specifically co-localized in nuclei of certain cells in HS epithelium. We also found that many of the PCNA positive hyperproliferative cells were also positive for c-Myc expression. These data demonstrate that 5’-cap□dependent translation is a potential pathway underlying the SCC pathogenesis in chronic HS lesions. Furthermore, being a druggable target, inhibition of eIF4F may block lesion-associated lethal SCCs in HS patients.

## Introduction

Hidradenitis suppurativa (HS) is a chronic, recurrent, inflammatory skin disease, which is often associated with painful and distressing symptoms ^1,2^. HS primarily affects apocrine-gland-rich regions of the body such as the axillary, inguinal, and anogenital areas. Young adults aged between 20 and 40 years have the highest incidence rates (0.05%~4%) and women are more frequently affected by HS compared to men ^3,4^. Patients with progressive disease develop inflammatory nodules, abscesses, and tunnels formation under the skin. HS disease severity is usually graded on the Hurley staging system, which can be classified into three stages (I, II, and III) ^5^. While several risk factors including inherited genetic mutations, hormonal status, obesity and smoking are known to be associated with disease severity in HS, the molecular drivers of HS are largely unknown ^6^. Until now, a surgical treatment remains the ultimate option for patients with Hurley stage II/III disease ^7,8^. Additionally, recent clinical reports of chronic HS demonstrate that epidermal hyperproliferation in inflammatory environment could lead to local spread of malignant lesions that often develop into squamous cell cancer (SCC) ^9–12^. Of the cutaneous malignancies, only the association of HS with SCC was statistically significant ^13^. However, the exact mechanism by which HS-associated epithelial hyperplasia is transformed into SCC remains unknown.

Increased protein synthesis with increased ribosome number, is important for cells to transition from quiescence to proliferation. This is also a critical step in malignant tumor progression ^14^. Translational control aberrantly dysregulates cell cycle and proliferation. eIF4E is the rate-limiting factor for 5’-cap□dependent translation initiation and thus an important rate-limiting regulator of protein translation initiation ^15^. eIF4E function is controlled the availability of 4E-BPs. The phosphorylation at Ser-209 mediated by the eIF4G-associated kinases the MAPK-interacting protein kinase (MNK)1 and MNK2 plays a decisive role in this regard ^16^. Formation of a ternary initiation factor complex involving eIF4E, eIF4G, and eIF4A1 is a key to provide the pathologic role of 5’-cap translation. We recently reported that increased expression of phosphorylated eIF4E, eIF4G, and eIF4A1 is associated with human and murine skin SCCs. We also found that enhanced eIF4E translation regulated targets, Cyclin D1 and c-MYC, are overexpressed in malignant skin lesions. Then, we showed that blockade of eIF4E signaling exerts potent antitumor effects in skin SCC, while MARK signaling inhibition attenuates eIF4E components association and SCC development in human xenograft murine model ^17^.

The eIF4F complex, consisting of the 5′ mRNA cap-binding subunit eIF4E, the scaffolding protein eIF4G, and the RNA helicase eIF4A, recruit mRNA to the 40S ribosome and regulates the cap-dependent mRNA translation process ^18^. The elevated expression and aberrant activity of this complex have been implicated in the etiology of a broad spectrum of human tumors. This involves selective translation of mRNAs needed for proliferation, metastasis, and multidrug resistance in tumor growth ^19–21^. Disruption of eIF4 complex formation either by blocking the eIF4E-eIF4G interaction or by targeting eIF4A, can cause cell cycle arrest followed by apoptosis induction in cancer cells ^18^. Importantly, the mitogen-activated protein kinase (MAPK) and mammalian target of rapamycin (mTOR) pathways play important role in the regulation expression of eIF4E components and their functional outcomes; and both of these pathways are involved in the pathogenesis of skin cancer ^22–24^.

In this study, we provide data to show that 5’-cap-dependent translation is augmented in HS lesional skin as we observed wide-spread presence of eIF4E/eIF4A1/eIF4G co-localization. We also demonstrate that two of eIF4F key translational targets, Cyclin D1 and c-MYC, are enhanced and co-expressed in specific cells within lesional epidermis. Moreover, eIF4F-associated RAS and PI3K oncogene signaling networks are highly enriched in HS transcriptome and their integration may contribute transcriptional amplification of c-MYC. Collectively, our work provides novel insights into the role of eIF4F complex in chronic severe HS disease and underscores a possible mechanism for oncogenic activity in HS epithelial cells. Inhibition of eIF4F would open new avenues for the development of therapeutic intervention of HS lesion-associated aggressive and lethal SCC.

## Methods

### Human subjects

The Institutional Review Boards of the University of Alabama at Birmingham approved the protocol (IRB-300005214) for obtaining surgically discarded skin tissues from heathy and HS (Hurley stage 2 or stage 3) subjects. Surgical excisions from 8 patients with HS (axillary and groin peri- and lesions) as well as 8 healthy controls from breast and tummy reduction surgery were collected. The aliquot of these tissue samples was stored at −80°C or fixed in 10% neutral buffered formalin until further study. In addition, a portion was used to prepare and culture epithelial cells from those tissue.

### Immunohistonchemistry (IHC)

Excised Tissues were fixed in 10% neutral buffered formalin for 24 hr at RT, then stored in 70% ethanol. Tissues were embedded in paraffin and tissues were cut into 5 μm sections using a microtome (HM 325, Thermo Fisher Scientific). This was followed by staining with hematoxylin and eosin (H&E). For this, slides were deparaffinized, rehydrated in a graded series of alcohol and microwave-treated in a citrate buffer (pH 6.0). For IHC, endogenous peroxidase activity was blocked using 0.3% hydrogen peroxide. After blocking, anti-eIF4E (1:100, Novus, Cat# NBP2-66802), anti-eIF4A1 (1:200, Cell Signal, Cat#2490), and anti-eIF4G (1: 100, Cell Signal, Cat#2469) were added to slides, which were incubated overnight at 4°C. On the following day, the corresponding HRP-linked secondary antibody was applied for 45 min at RT and then washed with PBS (3 times, 10 min). DAB solution was used to develop the signal. Sections were then visualized under KEYENCE BZ-X710 digital microscope (KEYENCE) and further analyzed with Bz X analyzer software (KEYENCE).

A semi-quantitative analysis was performed to evaluate IHC immunostaining as previously described ^25^. Briefly, the epidermal distribution of stained cells was assessed by the ratio of the area of positive cell layers over the total area of cell layers; for dermal analysis, the relative numbers of cells per square micrometer were determined by manually counting stained cells from eight random fields per sample.

### Confocal Immunofluorescence (IF) staining

For IF staining, skin sections were de-paraffinized, rehydrated and then incubated in antigen unmasking solution according to the manufacturer’s instructions (Vector laboratories). Sections were blocked in blocking buffer containing 5% normal goat serum in PBST (PBS+0.4% Triton X100) for 1h at 37□°C. Sections were then incubated with primary antibodies against proteins for anti-eIF4E (1:100, Novus, Cat# NBP2-66802), anti-eIF4A1 (1:100, Cell Signal, Cat#2490), anti-eIF4G (1: 100, Cell Signal, Cat#2469), anti-Cyclin D1 (1:200, Abcam, Cat# ab16663), anti-MYC (9B11) mouse mAb (1:200; Novus, Cat#2276), anti-PCNA (1:500; Abcam, Cat# ab264494), and anti-CDK4 (1:200; Cell Signal, Cat# D9G3E) in blocking solution for overnight at 4□°C. Sequential staining was done for visualization of more than one target in single specimen. After washing (3 times, 10 min each) with PBST, sections were re-incubated with indicated Alexa-Fluor□conjugated anti-goat or anti-rabbit secondary antibodies (1:200, Invitrogen). Sections were fixed in DAPI containing Vectashield antifade mounting medium (H-1200, Vectorlabs). Sections were then visualized under FLUOVIEW FV3000 confocal microscope (Olympus, USA) equipped with FV3000 Galvo scan unit and FV3IS-SW version 2.3.2.169 software.

### Isolation of primary keratinocytes and cells culture

HS and healthy skin tissues were cut into small pieces and were digested with 2.5U/ml Dispase-II (Roche, Cat#04942078001) in Hanks’ Balanced Salt Solution (HBSS, w/o Ca++ Mg++ (Corning, Cat#21-022-CM) overnight at 4°C. After digestion, the epidermis was peeled off and minced. This minced epidermis was incubated with 0.05% trypsin-EDTA at 37°C for 40 mins for further digestion. The suspension was then filtered through 40μm cell strainer and spun down at 300g at 4°C for 5 mins. Collected cells were counted at a density of 100,000 per well and seeded on matrigel pre-coated 6-well plates supplied with KBM-Gold keratinocyte growth medium (Lonza, Cat# 00192060) at 37°C in a humidified chamber with 5% carbon dioxide. We used keratinocytes 2-4 passages for performing experiments.

### Cell proliferation assay

Skin-derived keratinocytes were trypsinized, counted on a hemocytometer, and seeded on a white background plate (Thermo Scientific, Cat#354620) at a density of 5,000 per well in triplicate. These cells were allowed to grow further for 7 days at 37°C in a humidified chamber with 5% carbon dioxide. After 0,2,4 and 6 days, luminescent cell viability assay was conducted using CellTiter-Glo 2.0 kit (Promega, Cat# G9242) according to the manufacturer’s instructions. Fluorescence intensity was measured at 485–500nm_Ex_/520–530nm_Em_.

### qRT-PCR

Total RNAs were extracted with TRIzol reagent (Invitrogen, Cat# 15596026) and reverse-transcribed using the SuperScript III First-Strand Synthesis System (Invitrogen, Cat#18080051) with Oligo(dT)_20_ primers. For gene expression, qPCR was performed with Fast SYBR® Green Master Mix Real-Time PCR Master Mix (Applied Biosystems, Cat#4385612) on QuantStudio 12K Flex (Applied Biosystems) machine. The cycling acquisition program was as follows: 95□°C 20 sec, 40 cycles of 95□°C for 3□sec, 60□°C for 30 sec. We used human TaqMan probes from Thermal Fisher for Cyclin D1 (assay ID Hs00765553-m1) and c-Myc (assay ID Hs00153408-m1). Regular Primers used are list in the Supplementary Table1.

### Western-blot assay

Protein quantification and western blot analysis were performed as described previously (16). Briefly, whole-tissue lysates were prepared in RIPA buffer and sonicated. Tissue lysates were then centrifuged at 10,000g for 15 minutes at 4 °C; the supernatant obtained was used for protein concentration estimation by Bio-Rad DC protein assay kit (Bio-Rad, CA). Approximately 30-40 μg of protein was loaded into each well of an SDS-PAGE gel, electrophoresed, and transferred onto a polyvinylidene difluoride membrane. Membranes were blocked in 5% milk in Tris-buffered saline-Tween 20 for 1 hour and then incubated with primary antibodies for anti-p-eIF4E (Ser209) (1:1000, Abcam, Cat# ab76256), anti-eIF4E (1:1000, Novus, Cat# NBP2-66802), anti-eIF4A1 (1:1000, Cell Signal, Cat#2490), and anti-eIF4G (1: 100, Cell Signal, Cat#2469) overnight at 4°C. After three consecutive washes with PBST, membranes were incubated with horseradish peroxidase□conjugated secondary antibodies diluted in 5% milk for 2 hours, washed with PBST, and developed using enhanced chemiluminescence detection reagent. For sequential β-Actin antibody (1:20,000, Sigma-Aldrich #5316) re-probing, blots were stripped using Restore Plus Western Blot Stripping Buffer (Pierce Biotechnology) according to the manufacturer’s instructions. Band intensity was determined by densitometry and normalized to β-Actin.

### RNA-seq analysis

Published data was retrieved in FASTQ format from the following accession numbers: GSE155176 (whole tissue transcriptome of HS lesional, HS non-lesional, and healthy skin) ^26^ and GSE125380 (transcriptome of total mRNA in eIF4A-inhibitor-treated mouse pancreatic ductal adenocarcinoma organoids) ^27^. Sequencing reads were mapped to Gencode GRCh37.hg19 and Gencode GRCm38.p4, respectively, using STAR version 2.5.2b (options:– outReadsUnmapped Fastx;–outSAMtype BAM SortedByCoordinate;–outSAMAttributes All). Transcript abundances were calculated using Cufflinks version 2.2.1 with options–library-type fr-firststrand; -G; -L. Cuffmerge was then used to merge the transcript files from Cufflinks into one file. Following Cuffmerge, Cuffquant was used to quantify the transcript abundances, followed by differential gene expression using Cuffdiff. Differentially expressed genes with a P-value <0.01, as well as a log2fold change >1 were further analyzed further. We performed gene ontology (GO) analysis based on gene-GO association file (ftp://ftp.geneontology.org/go/gene-associations/gene_association.mgi.gz), and P-values were determined by Chi-squared test followed by Bonferroni correction.

### Gene set enrichment analysis (GSEA)

The list of 734 DEGs for eIF4F potential targets was used to analyze pathways by GSEA online tool (https://www.gsea-msigdb.org/gsea/index.jsp). We chose h.all.v7.3.symbols.gmt as the Gene sets database and Human_Gene_Symbol_with_Remapping_MSigDB.v7.4. chip as the chip platform.

### Statistical analysis

All data were statistically analyzed by GraphPad Prism 9.0. The data are expressed as the mean ± standard deviation (SD). Significant differences between groups were analyzed by two-tailed unpaired Student’s t test. P < 0.05 was considered significant.

## Results

### eIF4F complex components are highly expressed in HS skin

HS skin is characterized by epidermal hyperplasia and substantial immune cells infiltration in the dermis. We observed a thickened epithelium with epidermal psoriasis-like hyperplasia and massive infiltration of inflammatory cells in HS skin (Supplementary Figure 1A). To investigate potential association of eIF4 proteins with HS pathogenesis, skin biopsies were examined by IHC staining for assessment of the distribution and intensity (representative expression) of eIF4 major complex family proteins in healthy and disease skin. In healthy skin, slight to moderate expression for eIF4A1 and 4G was mainly observed in the basal layer of the epidermis; additionally, moderate expression of eIF4E protein was observed from the basal layer through the stratum spinosum towards the outer surface of the skin (Figure 1A, upper). This is consistant with our earlier observations ^17^. In contrast, enhanced expression of eIF4A1/4E/4G was seen in hyperplastic epidermis of HS skin, which expanded throughout the dermal compartment (Figure 1A, middle and bottom). Overall, these three proteins were more distributed in HS epidermis than in healthy skin (Figure 2B). Of note, in keratinocytes of both healthy and HS skin, eIF4A1 and eIF4G mainly expressed in cytoplasm; however, eIF4E protein localized to both cytoplasm and nucleus, which is consistant with its suggested role in exporting mRNA from nucleus to cytosol. Interestingly, the significant increase in abundance of three proteins was also observed in the dermal regions of HS skin compared to those in healthy controls, indicating the involvement of eIF4F proteins is not limited to epidermis but extended beyond to various cell types of the non-epidermal compartments. (Figure 1C, Supplementary Figure 1B). We also performed Western blot analysis to compare the expression of eIF4E and eIF4A1 in healthy and HS lesional skin samples. As shown in Supplementary Figure 1C and D, expression of those proteins was greatly elevated in lesional skin as compared to that in healthy skin. Taken together, eIF4 ternary complex might involve in the pathogenesis of HS, particularly driving its progression to late stage of the disease.

**Figure 1.**
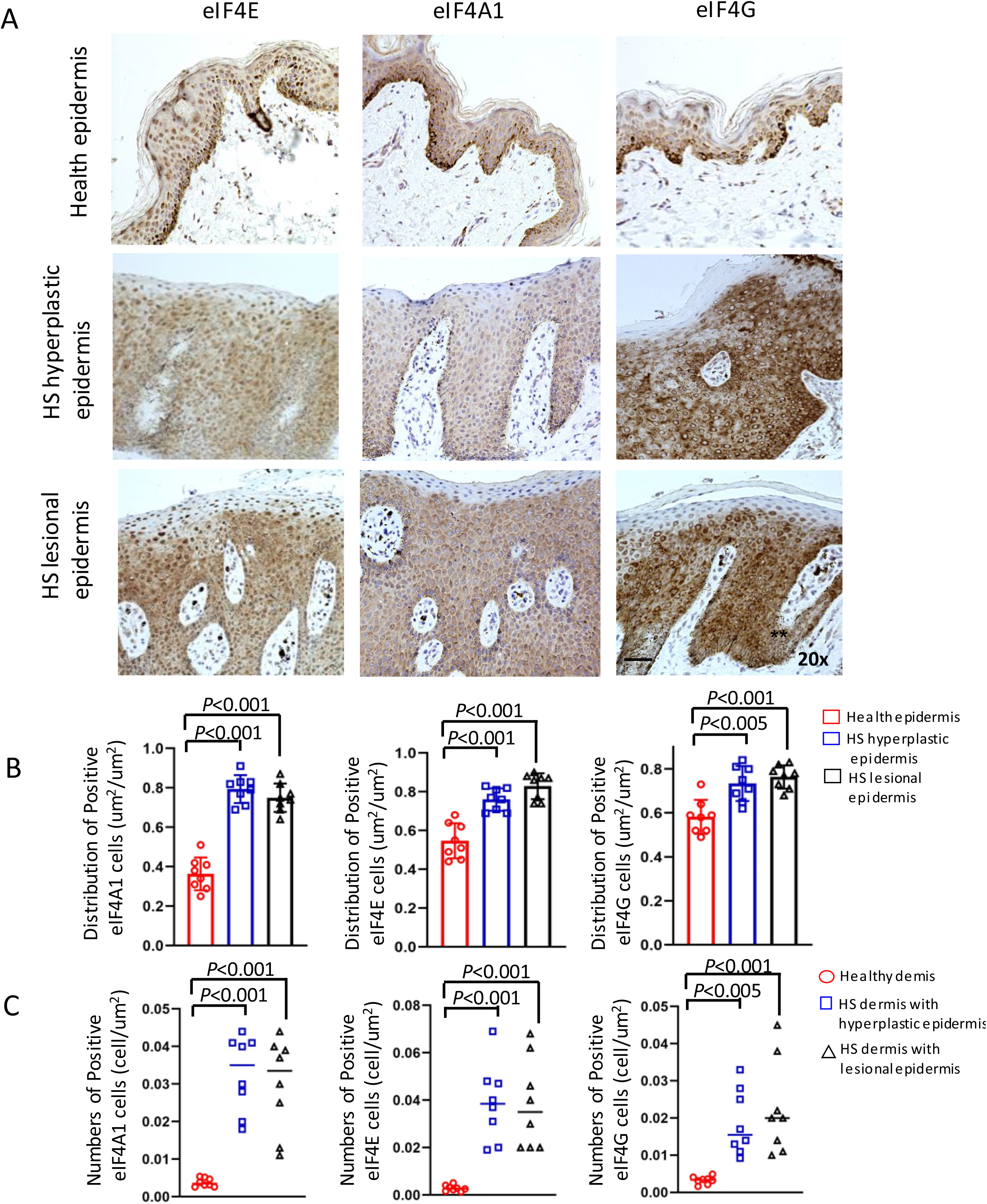
Induction in the expression of proteins associated with eIF4F complex was associated with HS severity (A) Representative images of human healthy and HS skin sections subjected to IHC staining for eIF4E, eIF4A1 and eIF4G expressions. Scale bar = 50μm. n=8 for healthy and HS skin, respectively. (B) Bar graphs showing the quantitative analysis for distribution of proteins such as eIF4E, eIF4A1 and eIF4G in per um^2^ area of healthy and HS epidermis (n=8 per group). (C) Graphs showing cell numeration of positive cells for eIF4F complex proteins in healthy and HS dermis (n=8 per group). Indicated P-value showing significance level of less severe hyperplastic epidermis and more severe HS lesions with respect to healthy controls. HS, hidradenitis suppurativa; IHC, immunohistochemistry.

**Figure 2.**
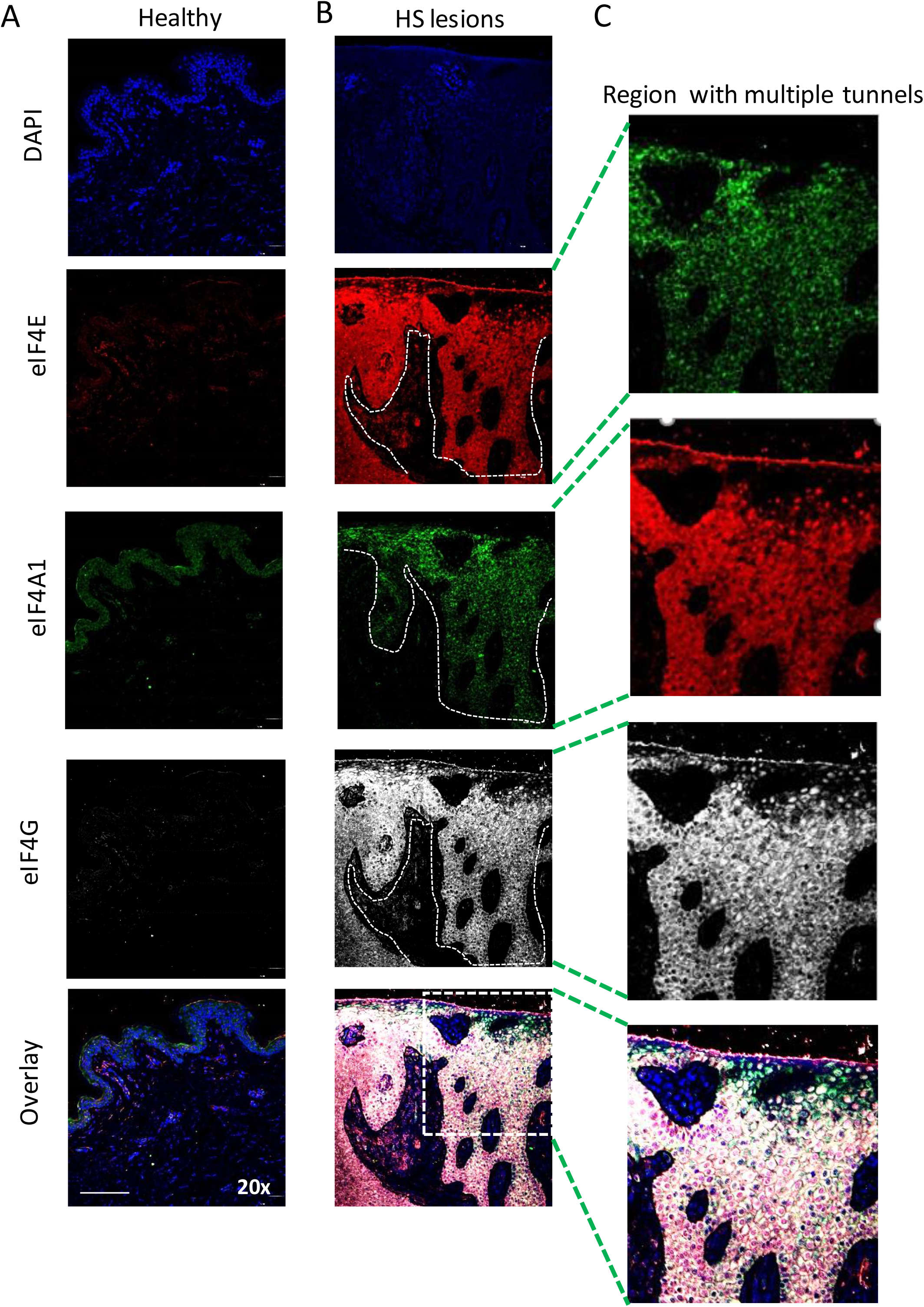
High-resolution confocal images showed the enhanced co-localizatinof eIF4F cmplex proteins in HS lesions relative healthy skin sections. (A and B) Representative images showing the enhanced expressioni of eIF4E (red), eIF4A1 (green), and eIF4G (grey) along with DAPI (blue) stained in nuclei in healthy human skin (A) and HS skin lesion (B). Slides were scanned using FLUOVIEW FV3000 confocal fluorescence equipped with FV3000 Galvo scan unit and FV3IS-SW software (Olympus, USA). White dot line indicating the junction of epidermal and dermal compartments in individual colored images while overlapping area of eIF4F complex proteins in overlay image. Scale bar = 100μm (C) Zoom area of images from (B). White dot box indicating the region with the overlapping expression for eIF4F three proteins. HS, hidradenitis suppurativa; IF, immunofluorescence.

### Subtissue anatomical localization of eIF4E/eIF4A1/eIF4G in HS skin

eIF4F complex has been previously reported to play conserved roles in translation initiation in the cytoplasm of eukaryotics ^28^. To explore the co-existance of different eIF4F components in HS keratinocytes, we utilized IF staining followed by confocal microscopy to capture high resolution images with respective to the subtissue and subcellular localization of these proteins. In healthy skin, weak signals for eIF4A1, 4E and eIF4G were confined mainly to the basal epidermal layer (Figure 2A, Supplementary 2A and B). However, in HS lesions, eIF4E and eIF4G and their co-localization were enhanced as ascertained by their mutually overlapping co-localizatin signals. These signals were widespread all over the hyperproliferative compartment, from the basal layer towards the suprabasal layers as indicated by the white dot lines (Figure 2B). The signal for eIF4A1 was mainly observed in the hyperproliferative suprabasal layers as well as in region withing multiple tunnels (Figure 2B, Supplementary Figure 2). In fact, three proteins show strong presence throughout regions of multiple tunnels as merged immunoreactivity demonstrated by bright yellow signals all over the tunnel forming cell (Figure 2C). eIF4A and eIF4G, positive staining was predominantly localized to cytoplasm, while immunofluorescence signal for eIF4E was identified in both the cytoplasm and nucleus, which is also consistent with our IHC results. Collectively, these results reveal that, in HS, eIF4F complex proteins assemble as a ternary complex within the cytoplasm of keratinocyte, which then aberrantly reprogram translation for genes involved in the differentiation and keratinization process.

### Potential target genes signature of eIF4F complex in HS lesions

Next, we sought to perform pairwise comparisons between the human HS transcriptome ^26^ and the core signature of eIF4A targets as established in mouse tumor organoids ^27^. There was substantial overlap in differentially expressed genes (DEGs), with a shared group of 734 genes that were differentially expressed in all three groups as indicated (Figure 3A, Supplementary Table-2). Among those, 516 and 218 were identified to up- or down-regulate, respectively (Figure 3B). So, this DEGs list serves as a reference dataset for eIF4F target identification in HS to point for the following analysis. We profiled the top 50 upregulated DEGs and the top 50 downregulated DEGs in Figure 3C. Compared with normal controls, previously identified genes were overexpressed in HS lesions, including those for known to be involved in tumorigenesis (*ACTRT3*, *CNN3*, and *PTPRN2*), skin inflammation (*IL10RA*, *CBARP* and *PTAFR*), keratinization process (*CCR2* and *FLl1*), skin fibrosis (*DOCK10*), innate immunity (*TLR7* and *BTK*) as well as keratinocyte growth and differentiation (*CRABP2*). In parallel, we enriched down-regulated genes for Wnt signaling (*WNT2B* and *WLS*), cell adhesion (*ELFN2*), epidermal homeostasis mediator (*KLK1* and *WDR72*), and inflammatory modulators (*TPH1* and *CXCL14*). Additionally, we validated the relative transcriptional abundances for some of interesting genes from DEGs list using RT-qPCR (Figure 3D). The results confirmed that *MYO1G* (associated with “B cell cytoskeleton rearrangements and antigene presentation”), *TBX2* (associated with “epithelial-mesenchymal transition and invasion in malignant cancer”), and *LEF1* (associated with “dermal fibroblast growth”) were significantly upregulated. While *CXCL14* (associated with “immune system normality”) and (associated with “transcriptional repression of TGFβ signaling pathway”) were decreased. Overall, transcriptome associated with normal skin was markedly dysregulated in HS skin and overexpression of eIF4F complex manifested strong negative effect on transcriptional products of skin-resident cells and their immune network.

**Figure 3.**
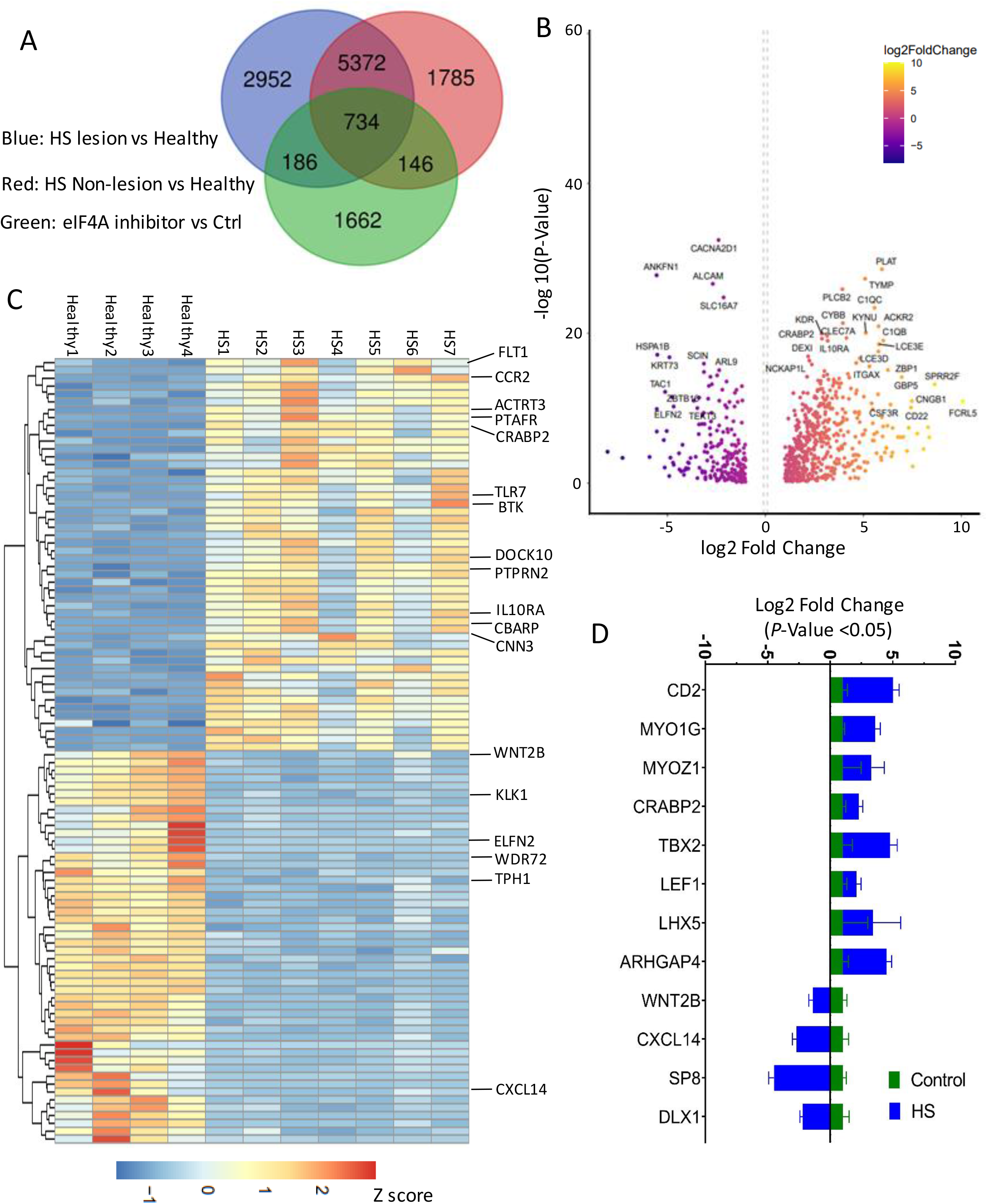
Target genes signature of eIF4F complex in HS lesions. (A) Venn diagram showing the intersection of eIF4F target genes between human healthy, HS lesion, and HS non-lesion transcriptome (GSE155176) and the core signature of eIF4A targets in mouse tumor organoids (GSE125380). The detailed number for overlapping genes derived from each comparison are shown inside the circles. (B) Volcano plot showing DEGs potentially regulated by eIF4F in HS. The X-axis represents the fold-change (log-scaled) and the Y-axis represents the *P*-value (log-scaled). Each symbol represents a different gene. Genes with –log10(FDR)□>=□20 are indicated. (C) Heat map representation of z-score and 2D clustering of DEGs generated from (A). Representative gene names are display on heat map. (D) qRT-PCR validation for selected genes from (A). Total RNA from healthy skin (n=4) and HS lesion skin (n=4) was extracted, reverse transcribed and analyzed. The Y-axis represents the fold-change (log-scaled). GAPDH expression was used for normalization. HS, hidradenitis suppurativa; DEGs, differential expression genes; FDR, false discovery rate.

### Activation of oncogenic signaling pathway in HS lesions

Next, gene ontology (GO) terms were assigned to the DEGs that were dysregulated in HS lesion compared to healthy counterparts, so as to identify their functions. GO terms successfully enrich several biological processes were significantly represented: immune system associated processes, such as T and B cell proliferation, inflammatory response, regulation of TNF production, and positive regulation of cell proliferation as well as PI3K signaling (Figure 4A). To further investigate how the products of these gene transcripts might act within HS progress, we performed relevant pathway enrichment analysis by mapping the results from 734 DEGs to the GSEA repository (Figure 4B). We found that the expression of genes involved in the inflammatory response and complement pathways were skewed toward HS disease progress. Specifically, interferon-gamma (IFN-γ) signaling was significantly upregulated in HS patients. Additionally, many of genes in RAS and PI3K signaling pathways were upregulated in HS. Key RAS/MEK/ERK and PI3K gene sets were subjected to an annotated hierarchical clustering procedure and presented as heatmaps. Among them, 33 genes in RAS/MEK/ERK pathway were upregulated in HS lesions, including those for cell growth (*KDR*, IGF1, and *PDGFRB*), NADPH oxidase complex (*NCF1* and *NOX4*) and inflammatory modulators *IL6* (Figure 4C). Fifteen genes were upregulated in PI3K pathway including immune-related hub gene *HCLS1* (Figure 4D). In sum, the augmented inflammation response-related pathways together with the activation of oncogene signaling networks may cause conducive environment for promoting tumorigenesis in severe HS stage II/III patients.

**Figure 4.**
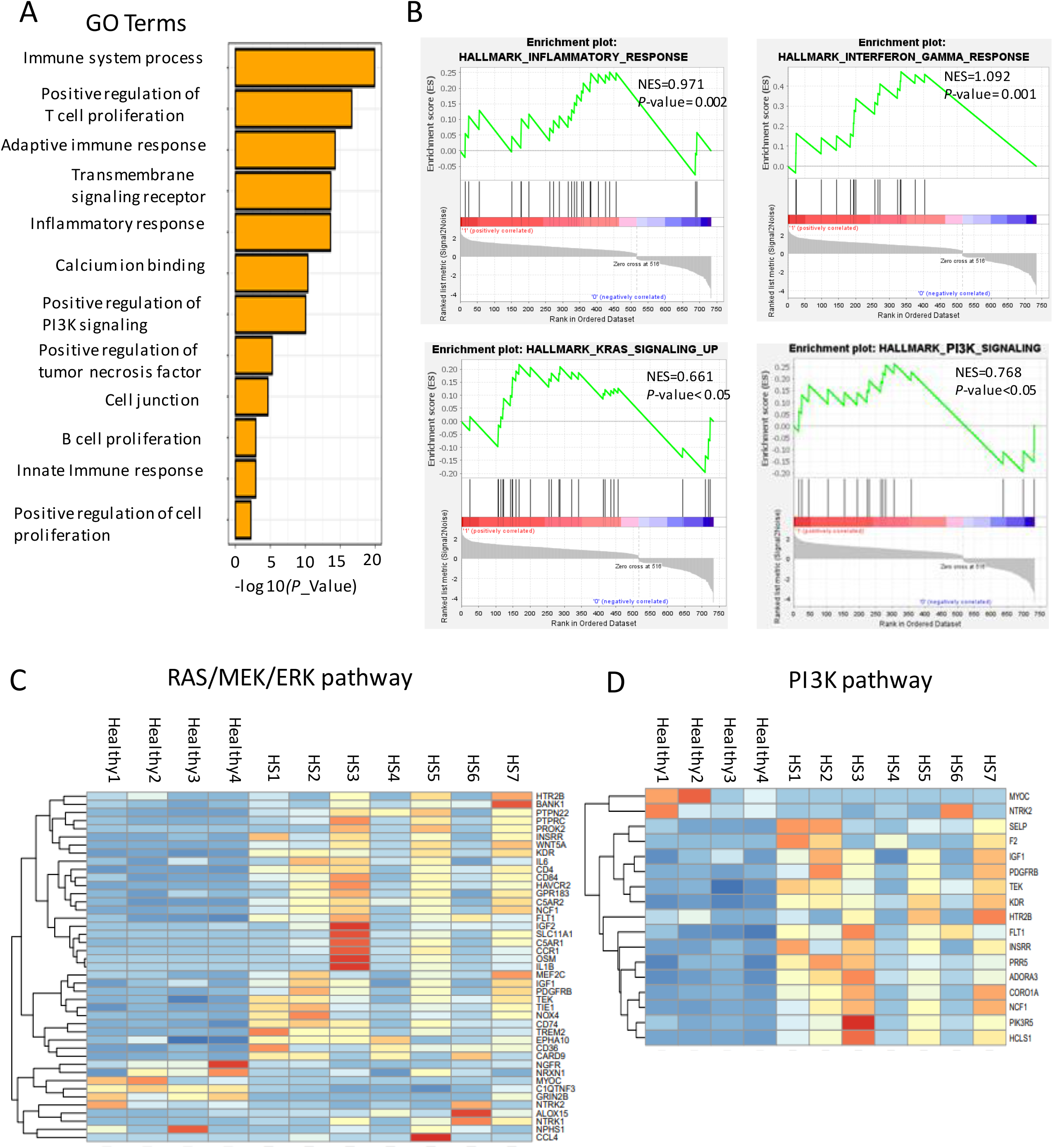
Activation of various signaling pathway in HS skin. (A) GO enrichment analysis showing top 12 ranked biological functions for eIF4F-related DEGs in human HS. The Y-axis represents the *P*-value (log-scaled). (B) GSEA tool was used to perform hallmark signaling as indicated for eIF4F reference gene sets according to Figure 3(A). (C and D) Heatmap showing the relative expression (z-scores) and 2D clustering of DEGs correlated with eIF4F in HS enriched in RAS/MEK/ERK pathway and PI3K pathway, respectively. Gene names are displayed on the heat map. GO, gene ontology; GSEA, gene set enrichment analysis.

### eIF4F complex is associated with enhanced HS skin cells

As RAS/MEK/ERK signals, crucial regulators of the cell processes such as cell proliferation, cell differentiation, and cell survival as well as the malignant transformation, were highly enriched in our reference dataset for eIF4F targets. Therefore, based on the dataset, we first identified two lists of all genes encoding proteins required for cell proliferation and cell cycle. We identified increased expression of 105 genes associated with the proliferation of several cell types. Transcription factors (*STAT4*, *PAX1*, and *LHX5*) for epithelia cell, tyrosine kinase-related gene *TEK* for endothelial cell, forkhead family member *FOXF1* for fibroblast cell, zinc finger protein *IKZF1* for T cell, transcription factor *LEF1* for B cell, and interleukin *IL12B* for nature killer cell was important to mentioned here (Figure 5B). We also found G1/S phase regulator *FBXL7* and positive regulator of cell-cycle progression *PROX1* exhibit high transcription pattern in HS samples (Figure 5C). Next, in our *ex vivo* validation, employing luminescence assay, we observed primary keratinocytes isolated from HS skin have enhanced proliferation capacity compared to those derived from healthy skin at the same time (Figure 5D).

**Figure 5.**
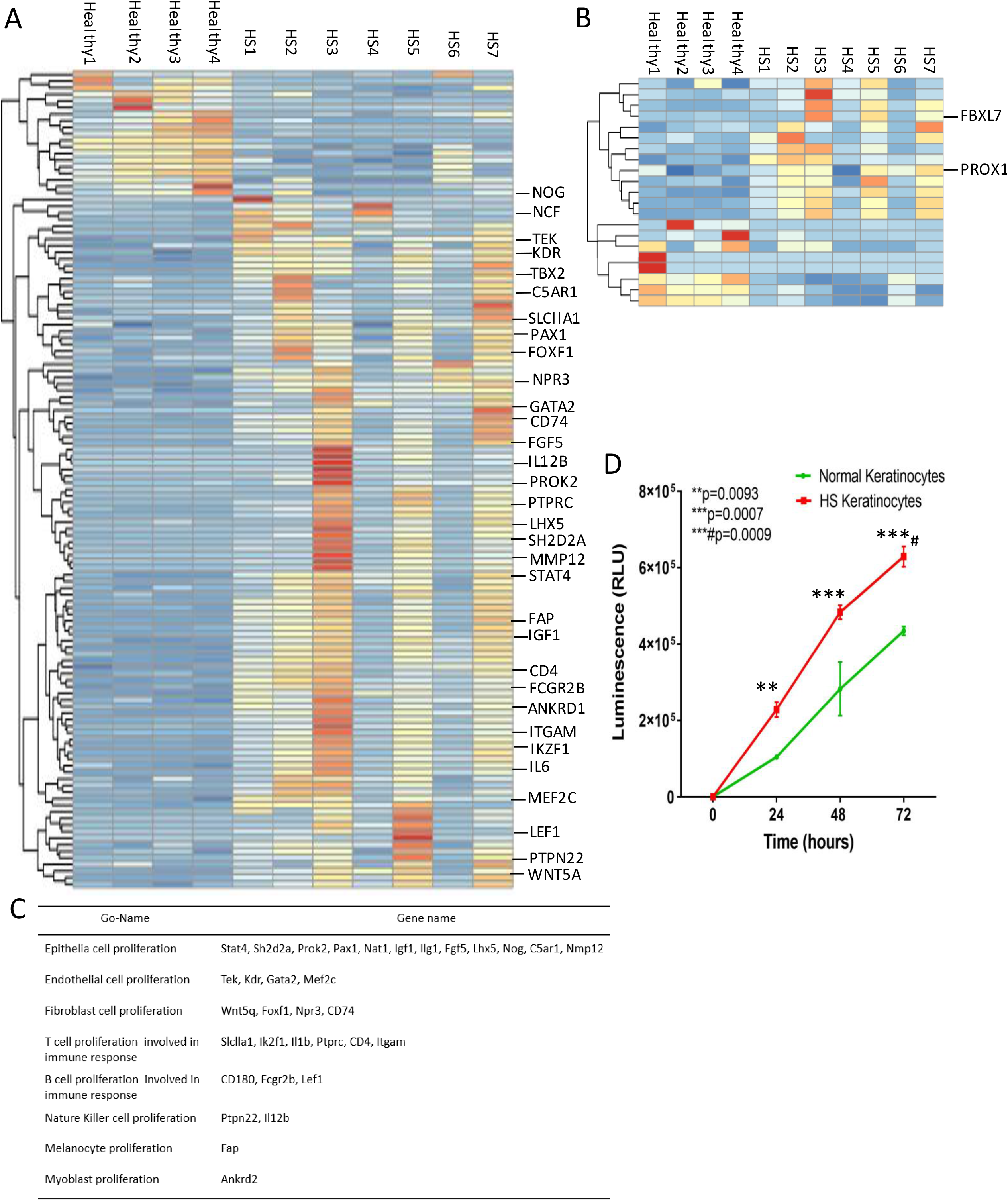
eIF4F-associated genes in HS lesions were involved in cellular proliferation. (A and B) Heat maps of eIF-4F-associated gene sets involved in GO term for ‘cell proliferation’ (A) and ‘cell cycle’ (B), respectively. The gene expression levels are standardized into z-scores and gene set is based on hierarchical clustering form. Representative genes are display on heat maps. (C) A list for selected key genes from (A) related in GO term ‘cell proliferation’ in various cell type. (D) Healthy- and HS-derived keratinocytes were seeded in triplicate into 96-well plates. The number of viable cells was measured over time using the CellTiter-Glo luminescent cell viability assay kit. One of three independent experiments is shown. Indicated *P*-value showing significance level compared with healthy control.

### Expression of nuclear Cyclin D1 and c-MYC is elevated in HS epidermis

We investigated the levels of two of eIF4 complex down-stream target proteins, Cyclin D1 and c-MYC. We observed that Cyclin D1 is accumulated in cytoplasm of cells in stratified epidermal layer in healthy skin (Supplementary Figure 6A, whiter arrow indicated), consistent with the previous observation ^29^. However, no expression for c-MYC was seen (Supplementary Figure 6A). In contrast, positive immunofluorescence signal of two proteins was observed in HS epidermis. In fact, cytoplasmic localization for Cyclin D1 was found in suprabasal cells of lesional epidermis (Figure 6B, right bottom panel; Supplementary Figure 3B), and a distinctive nuclear expression was mainly present in cells surrounding the multiple tunnels (Figure 6B, right upper panel; Supplementary Figure 3B). Additionally, nuclear localization of c-MYC, a feature for malignant cells, was strongly displayed throughout the lesional epidermis (Figure 6B, right panel; Supplementary 3B). Notably, strong co-localization signal for Cyclin D1 and c-MYC was found only in some epithelia cells forming tunnels. It is highly likely that this specific cell subpopulation may progress to neoplasm (Figure 6B, right bottom; Supplementary 3B). As a further confirmation at transcriptional level, we conduct qRT-PCR assay to exam their respective mRNA expression. Compared with healthy skin, upregulation of eIF4 proteins in HS skin coincides with significantly increased mRNA for *Cyclin D1* and *c-Myc* (Figure 6C)^30^. Further confocal microscopy confirmed that the active Cyclin D1/ CDK4 complex translocated into the nuclei in some of HS epidermal cells (Supplementary Figure 3C). We also observed that a hallmark of cancer cell proliferation PCNA, was up-regulated and co-localized with c-MYC in nuclei of hyperproliferative epidermal cells within HS lesions (Supplementary Figure 3D).

**Figure 6.**
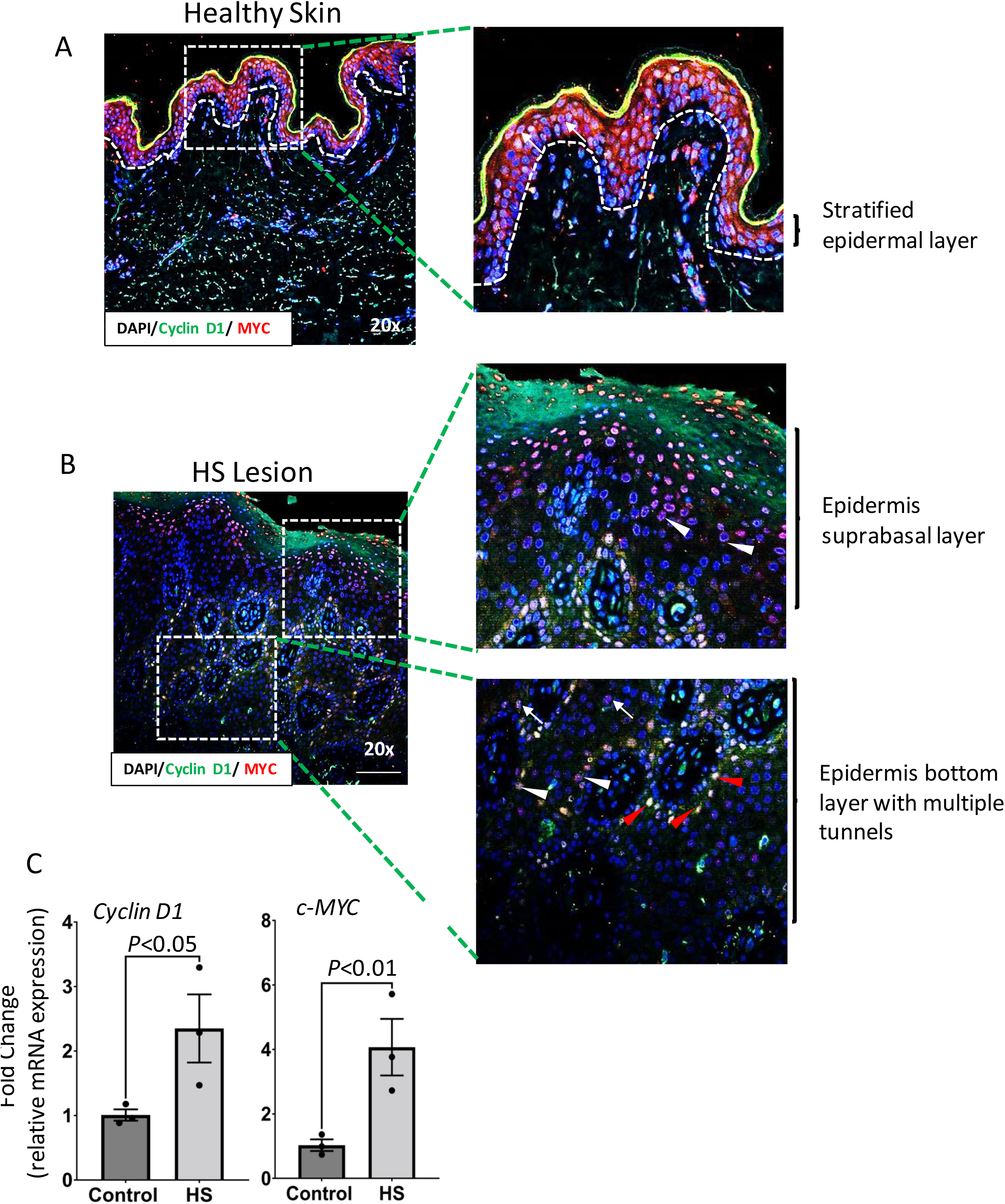
Overexpression patterns of Cyclin D1 and c-MYC in HS epidermis. (A) Representative confocal images of IF staining of Cyclin D1 (green) and c-MYC (red) along with DAPI (blue) in healthy human stratified epidermal layer. White arrow showing protein of Cyclin D1 localized in cellular cytoplasm of epidermal cells while expression of c-Myc was diffused in whole epithelia layer. (B) IF staining of Cyclin D1 (green) and c-MYC (red) along with DAPI (blue) in the lesional HS of human subjects. Magnified area of right upper panel showed that Cyclin D1 localized in celluler cytoplasm (white arrows) while e-Myc spefically expressing in cellular nuclei (white arrow head showing) in the suprabasal layer of HS epidermis. Right bottom panel showed the co-localization of Cyclin D1 and c-Myc inside nuclei at the basal layer of HS epidermis (red arrow). Slides were scanned using FLUOVIEW FV3000 confocal fluorescence equipped with FV3000 Galvo scan unit and FV3IS-SW software (Olympus, USA). Scale bar = 100μm. (C) Data from TaqMan qRT-PCR validate the enhanced expression of *Cyclin D1* and *c-MYC* mRNA in HS skin (n=3). The Y-axis represents the fold-change (log-scaled). GAPDH expression was used for normalization. HS, hidradenitis suppurativa; IF, immunofluorescence.

## Discussion

Overall, approximately 1%-3.2% of HS patients manifest SCC, which is more common in males (10). In familial HS, loss-of-function mutations have been reported in genes of the γ-secretase complex, including *NCSTN*, *PSEN1*, *PSENEN* and *PSTPIP1* ^31,32^. Among them, *NCSTN* mutants dysregulate keratinocyte proliferation and differentiation via activated Notch and PI3K signals ^33^, the latter of which has a prominent role in the pathogenesis of SCC ^34,35^. Moreover, *NCSTN*, *PSEN1*, *PSENEN* mutations have also been reported in sporadic HS from different cohorts of patients (for example, Caucasian, European, Asian) and the incidence rates are up to 3.2% ^36–39^. We also found a positive correlation between HS lesions and eIF4F-related induction of PI3K pathway (Figure 4B and D). However, to verify the relationship between Hurley stages II/III and *NCSTN* gene alterations using a large scale of patient samples, a genome-wide association studies (GWAS) study is necessary.

The metabolism of mRNA is a multiple step processing, including transcription, alternative splicing, and export of mRNA to the cytoplasm, which are time consuming process and therefore, cells use mature transcripts via translation by the energy efficient strategy for protein synthesis under various circumstances. In SCCs, the aberrant translation initiation rates dysregulate protein synthesis program ^40^. Increased of eIF4E/eIF4A1/eIF4G are associated with the enhanced survival and accelerated proliferation in SCC cells ^17,41,42^. Consistently, high abundance of eIF4F in the cytoplasm of both epidermis and dermis of HS biopsy (Figure 1 and Supplementary Figure1), may be associated with the possibility of promotion of lesional skin cells in tumor. Indeed, our observation of nuclear expressin of Cyclin D1 and MYC in some cells of HS epidermis layer (Figure 6B and Supplementary Figure 3B), point to their oncogenic role.

In addition, our observation that CDK4 is co-overexpressed with RAS in these cells (Supplementary Figure 3 C), further provide a mechanism underlying malignant progress of HS epithelial cells, which is consistent with the earlier demonstration of their role in human epidermal tumorigenesis ^43^.

In SCC progression from benign neoplasm, the maintenance of consistently high inflammatory microenvironment favors this transition ^44^. In HS, it is known that keratinocytes, dermal fibroblast and immune cells as well as skin resident DCs (LCs) and melanocytes, secrete a wide spectrum of immune-related clastogenic molecules (e.g., cytokines, chemokines, and growth factors), which create this inflammatory microenvironment. For example, it was known that IFN-γ generated by tumor-infiltrating cells enhances the survival of tumor cells and promote metastasis^45^. In this study, in addition to activated inflammatory pathways, interferon-gamma (IFN-γ) signaling is highly enriched in HS, which is also eIF4F-associated (Figure 4A and B).

Thus, our data provide a description of possible mechanisms by which 5’-cap translation initiation factors namely, eIF4E/eIF4A1/eIF4G proteins may be involved in highly conducive environment for malignant transformation and SCC development in chronic Hurley stage II/III patients. Furthermore, studies will provide a clear insight in this process.

## Acknowledgement

This work is supported by NIH grant RO1 ES026219 and NCI grant 5P01CA210946 and by intramural UAB funds to M.A.

## Declaration of no conflict of interest

Authors of this manuscript have no conflict of interest between them or anybody else regarding the scientific contents, financial matters, or otherwise.

---

**Supplementary Figure 1.**
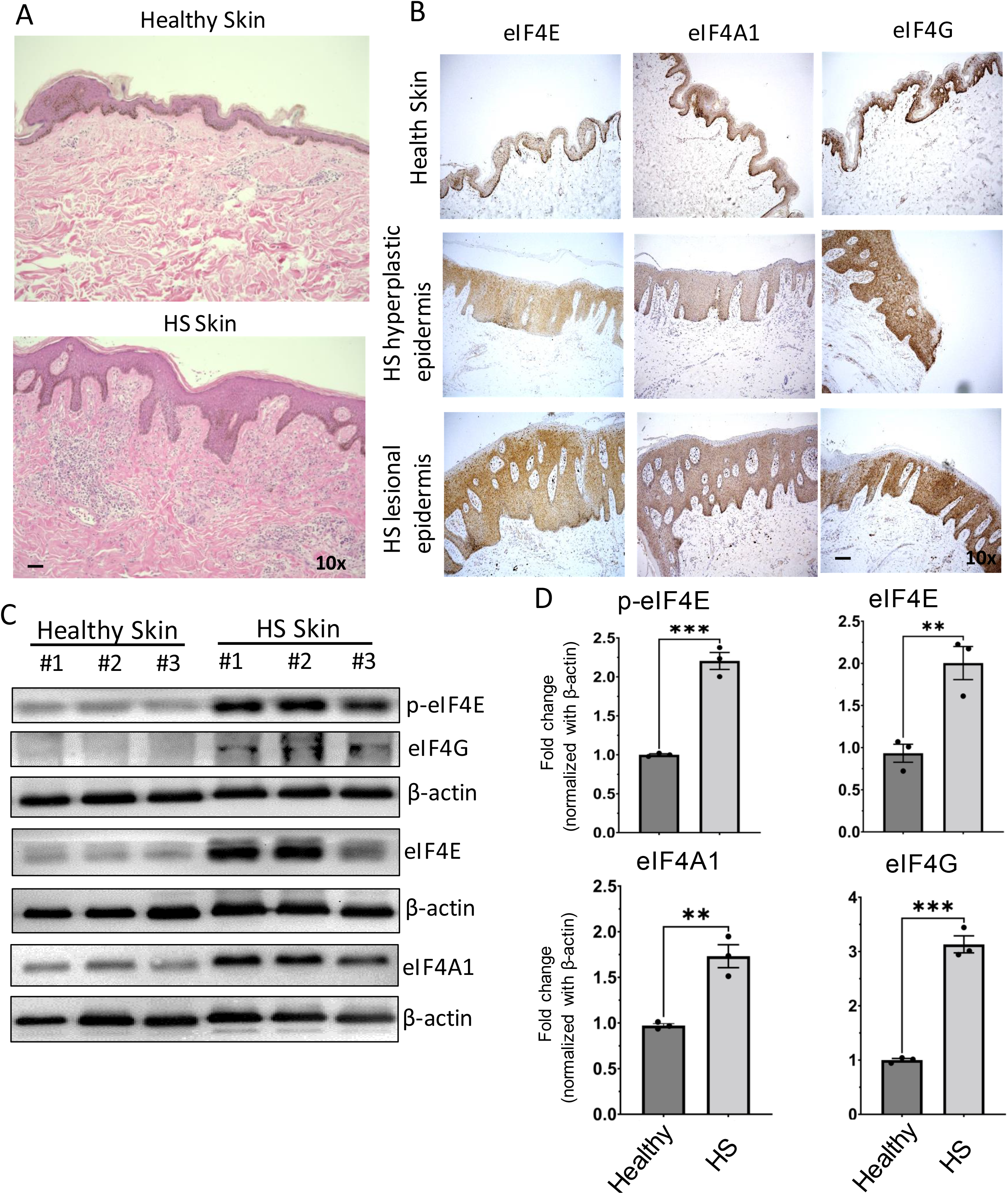
Expression analysis of eIF4E/eIF4A1/eIF4G in human HS. (A) Representative images of human healthy and HS skin sections subjected to H&E staining. Scale bar = 50μm. (B) IHC staining showing eIF4E/eIF4A1/eIF4G staining patterns in epidermis and dermis areas of human healthy and HS sections. Scale bar = 50μm. (C) Western blot analysis of eIF4E and eIF4A1 expression in three individual samples randomly selected from human healthy and HS skin group. β-actin was used as an endogenous control. (D) (D) Western blot analysis of p-eIF4E, eIF4E, eIF4A1, and eIF4G expression in human HS skin tissue compared with that in healthy skin tissue. Histogram showing densitometry analysis of band intensity and presented in fold change. Indicated *P*-value showing significance level compared with healthy control (n=3 per group). HS, hidradenitis suppurativa; H&E, hematoxylin and eosin; IHC, immunohistochemistry.

**Supplementary Figure 2.**
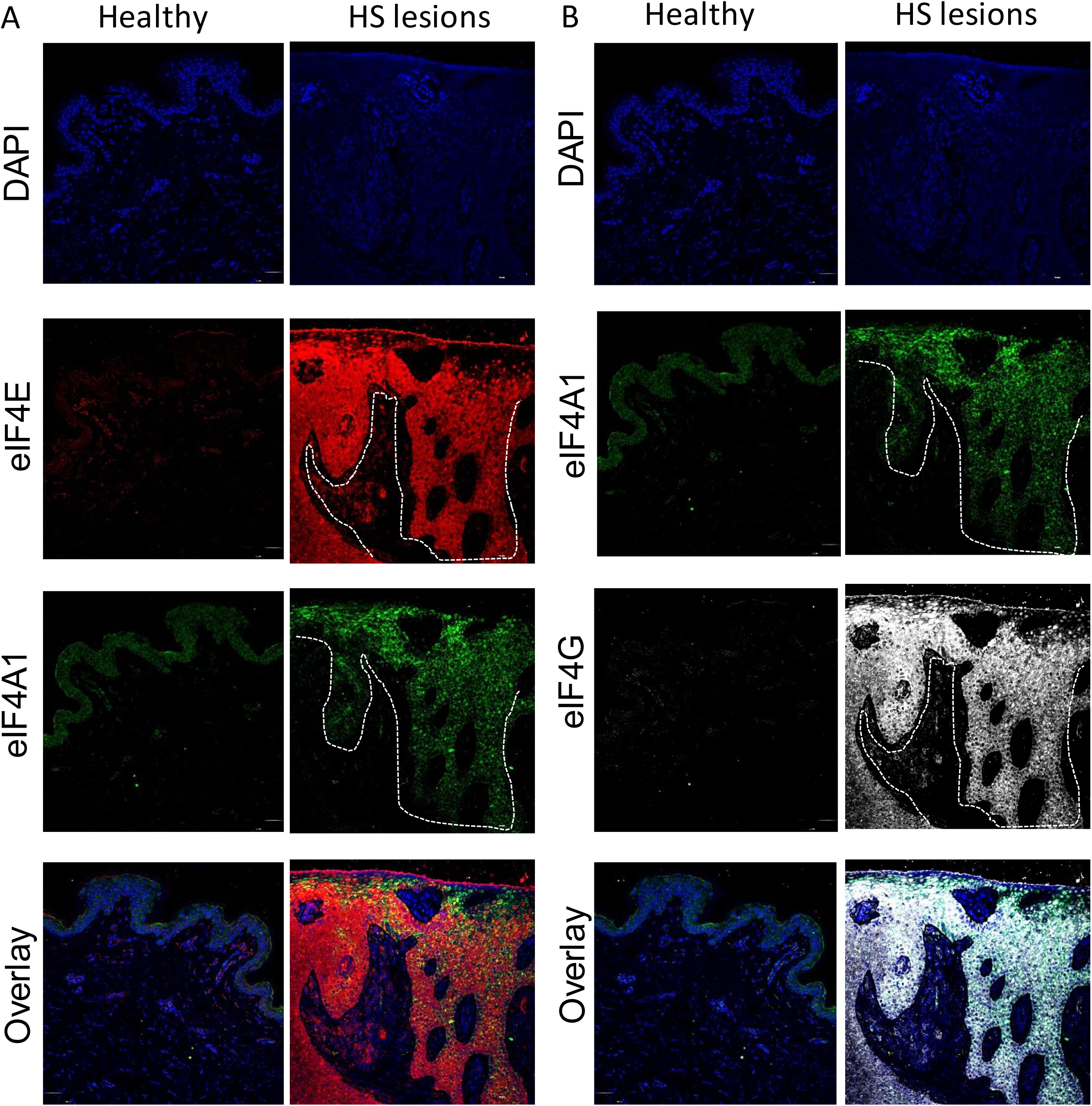
Mutual localization of eIF4E/eIF4A1/eIF4G components in human HS lesions. (A and B) Representative individual images of IF staining of eIF4E (red), eIF4A1 (green), and eIF4G (grey) along with DAPI (blue) in (A) healthy human skin and (B) HS lesion. Slides were examined under confocal fluorescence microscope (FLUOVIEW FV3000, Olympus, USA). White dot line indicating the positive IF staining area for individual eIF4E/eIF4A/eIF4G protein. White dot box indicating the region with the mutual overlapping expression for eIF4E/4G/A1. Scale bar = 100μm. HS, hidradenitis suppurativa; IF, immunofluorescence.

**Supplementary Figure 3.**
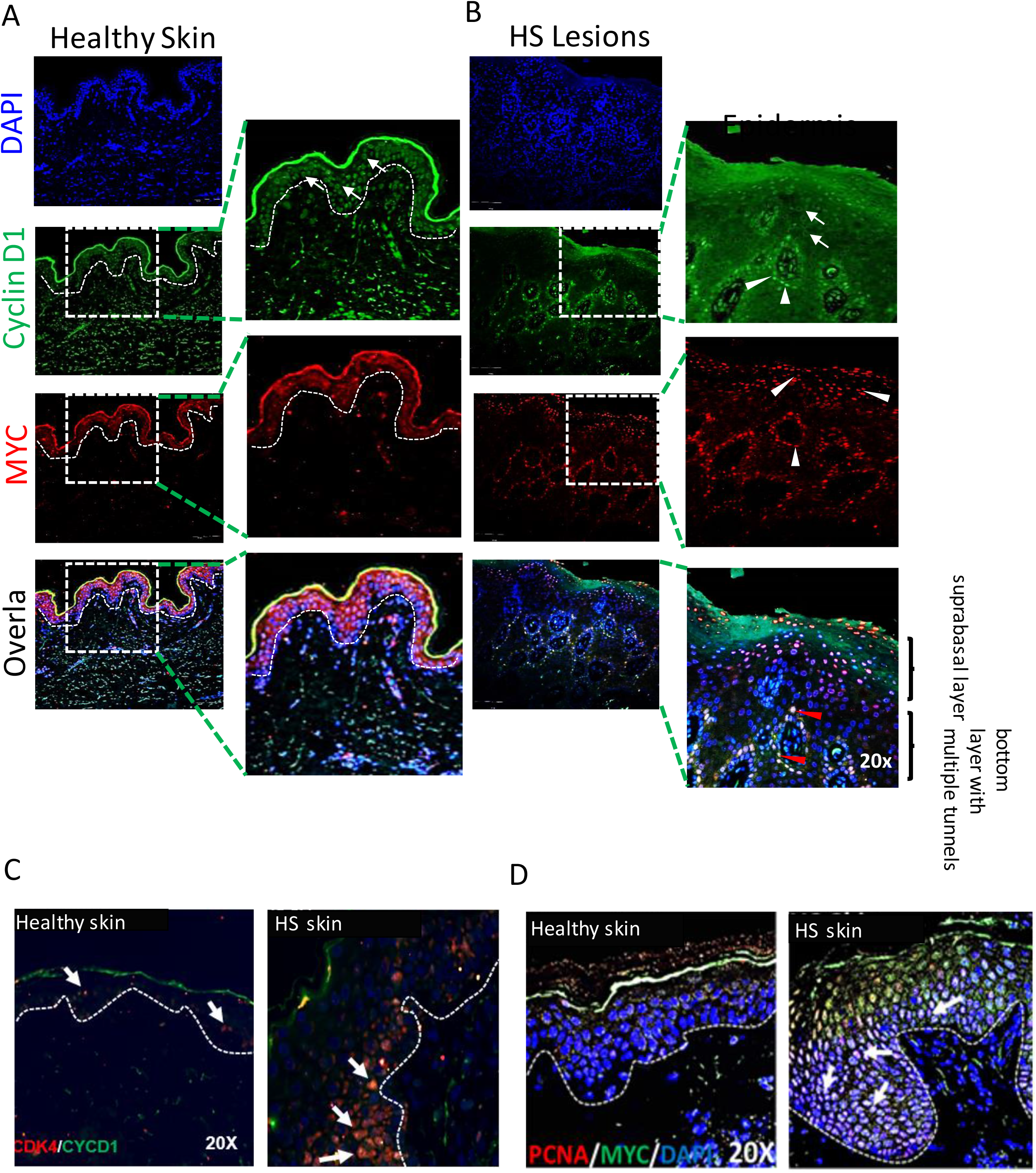
High expression of Cyclin D1 and c-MYC in HS epidermis. (A) Representative images of IF staining of Cyclin D1 (green) and c-MYC (red) along with DAPI (blue) in healthy human stratified epidermal layer. White dot line indicating the border between epidermis and dermis. White dot box indicating the magnification of area. (B) Representative images of IF staining of Cyclin D1 (green) and c-MYC (red) along with DAPI (blue) in human HS lesional epidermis. White dot box indicating the magnification of area. White arrow showing protein of Cyclin D1 localized in cellular cytoplasm. White arrow head showing protein of c-MYC specifically expressing in cellular nuclei. Red arrow head showing Cyclin D1 and c-MYC co-localized in nuclei. (C and D) Confocal images demonstrating the co-localization of Cyclin D1/CD4 and PCNA/MYC in hyperplastic epidermis of healthy and HS skin. White arrow showed the co-localization of respective proteins inside nuclei in HS epidermis. Slides were scanned using FLUOVIEW FV3000 confocal fluorescence equipped with FV3000 Galvo scan unit and FV3IS-SW software (Olympus, USA). Scale bar = 100μm. HS, hidradenitis suppurativa; IF, immunofluorescence.

