## Supplementary Table1 for "Dysregulated Protein Translation Control in Hidradenitis Suppurativa: Implication for Lesion-associated Squamous Cell Carcinoma Development"

### Supplementary table-1:

List of human primers:

| GENES | | FORWARD | REVERSE |
|---|---|---|---|
| CRABP1 | | 5' CAGGACGGGGATCAGTTCTA 3' | 5' CTGCAAAGGGAGACACAGGT 3' |
| CD2 | | 5' CCTTGGGTCAGGACATCAAC 3' | 5' TCATCGGTCTTCAGATGCTT 3' |
| Tbx2 | | 5' CTCCTTCTTCCCGGCACT 3' | 5' ACTGGTCCCACAGCTCCTT 3' |
| Arhgap4 | | 5' GCTGAGGTGGAGCTGGAATA 3' | 5' CCTCTGCAATGTGACTCAGG 3' |
| Myo1g | | 5' ACCAAGTGACCATGGAGGAC 3' | 5' TTGGCCACAGCATAGAGATG 3' |
| Lef1 | | 5' GACGAGATGATCCCCTTCAA 3' | 5' GGTAGGGCTCCTGAGAGGTT 3' |
| MYOZ1 | | 5' CTTGAACCTGGGCAAAAAGA 3' | 5' CCGTTGCTCTTGCTGTATGA 3' |
| WNT2B | | 5' TAGGTCTTGCCTGCCTTCTG 3' | 5' GCTGACACTCTCGGATCCAT 3' |
| LHX5 | | 5'AGTGCAAAACCAACCTCTCG3' | 5'TGCAGGTGAAACAGTTGAGG3' |
| CXCL14 | | 5'AAGCTGGAAATGAAGCCAAA3' | 5'TTCCAGGCGTTGTACCACTT3' |
| SP8 | | 5'AGGTTGGGATCGACTCCTCT3' | 5'ACTGGAACCCACTACGTTGC3' |
| DLX1 | | 5'CAAGGCGGTGTTTATGGAGT3' | 5'TGCTGACCGAGTTGACGTAG3' |
